# Identification of the gene coding for seed cotyledon albumin SCA in the pea (*Pisum* L.) genome

**DOI:** 10.1101/2022.03.12.484058

**Authors:** Anatoliy V. Mglinets, Vera S. Bogdanova, Oleg E. Kosterin

**Author notes:** Corresponding author; ORCiD 0000-0001-5955-4057.

## Abstract

Albumins SCA and SAA are short, highly hydrohylic proteins accumulated in large quantity in the cotyledons and seed axes, respectively, of a dry pea (*Pisum sativum* L.) seed. SCA was earlier shown to have two allelic variants differing in mobility in polyacrylamide gel electrophoresis in acid medium. Using them the corresponding gene *SCA* was mapped on Linkage Group V. This gene was used as a useful genetic and phylogeographical marker that still required electrophoretic analysis of the protein while the DNA sequence of the corresponding *SCA* gene remained unknown. Based the length, the positive charge in acid conditions and the number of lysine residues of SCA and SAA albumins, estimated earlier electrophoretically, the high throughput sequencing data available in public databases were searched for the respective candidates for the corresponding genes *SCA* and *SAA* genes among coding sequences residing in the region of the pea genome corresponding to the map position of *SCA*. Then we sequenced them in a number of pea accessions. Concordance of the earlier electrophoretic data and sequence variation indicated the sequence *Psat0s797g0160* of the reference pea genome to be the *SCA* gene. The sequence *Psat0s797g0240* could encode a minor related albumin SA-a2, while a candidate gene for albumin SAA is still missing (as well as electrophoretic variation of both latter albumins). DNA amplification using original primers SCA1_3f and SCA1_3r from genomic DNA and restriction by endonuclease *Hind* II made SCA a convenient CAPS marker in peas. Thus, the gene encoding the highly hydrophilic albumin SCA accumulated in pea seeds, with its alleles useful classification of pea wild relatives, is now identified in the pea genome and a convenient CAPS marker is worked out on its base.

## Introduction

Mature pea (*Pisum sativum* L.) seeds contain a number of albumins extractable from their mill with 5% perchloric acid, which were characterised in detail by Smirnova et al. (1990; 1992). A special attention was drawn by the two most abundant of these albumins, which are biochemically and immunologically related and quite short (about 100 amino acid residues) highly hydrophylic peptides. One of them predominates in the cotyledons and the other in the seed axis, both accumulating during seed formation and depleting while germination. They were respectively named SCA (seed cotyledon protein) and SAA (seed axis protein). Their gross amino acid content and the accumulation pattern in seeds left no doubt in their participation in substitution of water in dry seed cells, that is in a dehydrin-like function, although they differed in many respects from the known dehydrins and had much smaller molecules (Smirnova et al., 1992). While the SAA protein was electrophoretically monomorphic in peas, two allelic variants of SCA were revealed differing in electrophoretic mobility in 15% polyacrilamide gels containing acetic acid and urea according to Panyim & Chalkley (1969), which allowed to genetically map the relevant gene *SCA* (Smirnova et al., 1992; Rozov et al., 1993; Gorel et al., 1998) on linkage group V (corresponding to chromosome 3; see Smýkal et al. 2012; Kreplak et al., 2019).

Smirnova et al., (1992) found out that the fast electromorph SCA^f^ was frequent in the wild subspecies of the common pea (*P. sativum* subsp. *elatius* Aschers. et Graebn.) but was extremely rare in the cultivated subspecies *P. sativum* L. subsp. *sativum*. (Here the inclusive taxonomic system of peas according to Maxted and Ambrose (2001) is followed.) Kosterin and Bogdanova (2008) and Kosterin et al. (2010) noticed a strong concordance of the occurrence of SCA^f^ with that of the plastidic *rbcL* allele containing a recognition site for the *Hsp* AI restriction endonuclease, and a less strong concordance with that of the mitochondrial *cox1* allele containing recognition site for the *Psi* I restriction endonuclease. This concordance was interpreted in terms of the common phyletic origin and as evidence of existence of two different wild pea lineages. Based on this, different combinations (A, B and C) of alleles of the three mentioned dimorphic marker genes *SCA, rbcL* and *cox1* from different cellular genomes, respectively nuclear, plastidic and mitochondrial, were proposed for a simple classification of evolutionary lineages of the wild pea subspecies *P. sativum* subsp. *elatius* (Kosterin, Bogdanova, 2008; Kosterin et al., 2010), which was then used repeatedly (Zaytseva et al., 2012; 2015; 2018; Kosterin, Bogdanova 2021, Bogdanova et al., 2021 etc.). So the electromorphs of SCA appeared to be useful in the studies of genetic diversity of the pea crop wild relatives, which are important for involvement of their potentially useful genetic resources into breeding (Ali et al., 1994; Maxted, Kell, 2009; Coyne et al., 2011; Ford-Lloyd et al., 2011; Maxted et al., 2012). However, while the plastidic and mitochomdrial markers were scored by the CAPS approach involving DNA amplification and restriction, the *SCA* gene sequence remained unknown and analysis of this marker required more laborious protein electrophoresis. Molecular identification of this gene would be desirable to facilitate the analysis. This became possible when the pea nuclear genome was published by Kreplak et al. (2019). This communication is devoted to identification of the *SCA* gene in the pea genome, its brief characterisation and working out a convenient CAPS marker based on this gene.

## Material and methods

The gene SCA was sequenced from samples of DNA extracted in the course of our previous work (Kosterin, Bogdanova, 2008 to be consulted for details), from the following pea germplasm accessions: 721, CE1, CE11, JI1794, L100, PI344538, Pse001, Pe013, P015, P017, VIR320’ (*P. sativum* subsp. *elatius*), VIR3429, VIR4911, VIR5416, VIR7335, WL1238, Cameor (*P. sativum* subsp. *sativum*), VIR4871 (*Pisum sativum* subsp. *transcaucasicum* Govorov), VIR2759 (*Pisum abyssinicum* A. Br.), and WL2140 (*P. fulvum* Sibth. et Smith). Polymerase chain reaction to amplify the fragment corresponding to *Psat0s797g0160* was carried out in 20 μl of the PCR reaction mixture under a mineral oil layer with the following cycling parameters: 95°C for 3 min; 45 cycles consisting of denaturation at 94°C for 30 s, annealing at 56°C for 25 s and elongation at 72°C for 40 s; final elongation at 72°C for 5 min; for amplification of the fragment corresponding to *Psat0s797g0240* the same parametres were used but annealing was at 59°C. For restriction analysis, 5 μl of the resulting reaction mixture were digested with 1 unit of *Hind*II endonuclease according to manufacturer’s instructions and the products analysed on 1.5% agarose gel in TAE buffer. For sequencing, PCR products were purified by 20% polyethylene glycol 800 in 2.5 M NaCl. Sanger reaction was carried out using BrightDye Terminator version 3-100 (Nimagen, Netherlands) with the conditions recommended by the manufacturer for 50 cycles. The Sanger reaction products were purified using Sephadex G-75. Sequencing was carried out in Genomic Core Facility SB RAS, Novosibirsk. The sequences obtained in this study were submitted to European Nucleotide Archive with the following entry numbers OU953856-OU953865, OU953869-OU953881 for *SCA* and OU953866-OU953868, OU953882-OU953894 for alleles of *Psat0s797g0240*.

## Results and Discussion

### Candidate gene for *SCA* seed albumin in public databases

The *SCA* gene is mapped on likage group V (LGV) between the loci *His1* (coding for histone H1 subtype 1) and *coch* (Gorel et al., 1998) (its earlier published position behind *coch* (Rozov et al., 1993) was tentative as based on non-additive data with respect to these three loci). The potential candidates were searched in the annotated genome of *Medicago truncatula* Gaertn. making use of its syntheny with the genome of *Pisum* (Kalo et al., 2004). The bordering markers *coch* and *His1* of pea correspond to the loci with Gene IDs 11417633 and 25499208, respectively, therefore suitable candidates found in *Medicago truncatula* should map to chromosome 7, synthenic to LGV of *P. sativum* (Kalo et al., 2004, Kreplak et al., 2019) at physical position between 42,203,622 (GeneID: 11417633) and 45,488,994 (GeneID: 25499208) on NC_053048.1 (*M. truncatula* strain A17 chromosome 7). This region contained 421 coding sequences, of which three neighbouring loci were annotated as “18 kDa seed maturation protein”. Two of them, *LOC11421661* and *LOC11437338*, encoded polypeptides of 105 and 101 amino acids, respectively. The third locus, *LOC11437936* encoded polypeptide of 177 amino acids. These polypeptides were used as a query to search the *P. satvum* genome assembly at https://urgi.versailles.inra.fr/blast. All three searches retrieved the same hits, *Psat0s797g0240* and *Psat0s797g0160*, separated by about 25 Kb on the scaffold 00797 not attributed to any chromosome and *Psat3g068920* on chr3LG5 with physical position between the loci *coch* and *His1*. *Psat3g068920* encoded polypeptide of 190 amino acids and probably corresponded to the *LOC11437936* of *M. truncatula* while Psat0s797g0240 and Psat0s797g0160 probably corresponding to LOC11421661 and LOC11437338 both encoded polypeptides of 101 amino acids and were concluded to be ideal candidates to represent the SCA locus.

*Psat0s797g0160* with position 57,48857,939 on scaffold00797 annotated as “Late embryogenesis abundant (LEA) group 1” encoded a polypeptide of 101 amino acid residues including 18 positively charged residues, of which 10 were lysines. *Psat0s797g0240* with position 83,88584,190 on scaffold00797 also annotated as “Late embryogenesis abundant (LEA) group 1” encoded a polypeptide of 101 residues including 20 charged residues of which 11 were lysines. The earlier obtained data on amino acid composition of the SCA protein were as follows. The slow electromorh SCA^s^, common in cultivated peas, was estimated by the incomplete succinylation method to possess 17 positively charged residues, including 9 lysines, and, together with data on gross amino acid content, as being ca 107 residues long (Smirnova et al., 1992). This was rather a good correspondence of data from protein chemistry and sequencing. The estimate of 9 rather than 10 lysine residues in SCA^s^ with the incomplete succynilation method (Smirnova et al., 1992) may be explained by a tandem of two lysine residues in positions 47-48 of the deduced protein product, which could not both bound to succinic acid residues for steric reasons.

Hence the size and amino acid content of the protein products and genome location of both *Psat0s797g0160* and *Psat0s797g0240* corresponded to SCA (Smirnova et al., 1992; Gorel et al., 1998). To choose between them for a candidate for SCA we made use of the genetic data. SCA was shown to be dimorphic with allelic variants differing in respect of the number of positively charged amino acid residues while SAA was monomorphic in this respect (Smirnova et al., 1992). Sequences available in sequence read archives (SRA) at NCBI containing data of high throughput resequencing of pea accessions were used to assemble alleles of *Psat0s797g0160* and *Psat0s797g0240* from pea accessions W6_2107 (BioProject PRJNA431567), JI1794, WL 2140 (*Pisum fulvum* Sibth. et Smith) (BioProject PRJNA431567), JI2202 (*Pisum abyssinicum* A.Br.) (BioProject PRJNA285605), 711 (BioProject PRJEB30482), 721 (BioProject PRJNA431567, PRJEB30482). Five of these accessions were involved in our previous electrophoretic studies of SCA (Kosterin, Bogdanova, 2008); Cameor and JI1794 were shown to have the slow electromorph SCA^s^ while WL2140, 711 and 721 had the fast electromoph SCA^f^. There was some sequence variability among alleles of Psat0s797g0160. However, two nucleotide substitutions differed in the alleles of Cameor, W6_2107 and JI1794 from those of WL2140, JI2202, 711, 721, namely T/G in the position 215 (from start codon) and A/C in the position 238. The T/G substitution changed valine for glycine, and the A/C substitution changed asparagine to histidine, wich is positively charged under conditions of acetic acid-urea PAAG used for SCA analysis. Thus, the latter amino acid replacement affects electrophoretic mobility of the encoded protein and is associated with fast electromorph. Allelic variants of Psat0s797g0240 did not carry amino acid substitutions associated with the change of electrophoretic mobility. This allowed us to nominate Psat0s797g0160 as a candidate for the SCA gene.

### Concordance of the sequence variation of the candidate gene for *SCA* with SCA electrophoretic pattern

To confirm *Psat0s797g0160* to be the *SCA* gene we resequenced it in 20 pea accessions in which SCA was previously studied electrophoretically (Smirnova et al., 1992; Kosterin, Bogdanova, 2008; Kosterin et al., 2010). To design primers matching 3’ and 5’ non-coding regions we searched public databases for pea sequences coding for the same protein as *Psat0s797g0160*. The search retrieved the sequence PUCA013656022.1 (*Pisum sativum*, cultivar Gradus No 2 flattened_line_3009, whole genome shotgun sequence) containing the coding sequence as well as 5’ and 3’ non-coding regions. The primers Ps_SCA1_3F (5’ GCATCATACTCTTCAACACAT) and Ps_SCA1_3R (5’ GTAGGAACATTCACAACATCA) were designed to match those non-coding regions. We sequenced the coding sequence of the *SCA* gene in two groups of 10 pea accessions each: (i) L100, 721, VIR320’, CE11, PI344538, Pe013 (*P. sativum* subsp. *elatius*), VIR2759 (*Pisum abyssinicum* A.B.), VIR3429, VIR7335 (*P. sativum* subsp. *sativum*) and WL2140 (*P. fulvum*) which were shown to have the fast SCA electromorph (SCA^f^), frequent in wild peas, and (ii) accessions CE1, JI1794, Pse001, P015, P017 (*P. sativum* subsp. *elatius*), VIR4871 (*Pisum sativum* subsp. *transcaucasicum*), VIR4911, VIR5416, WL1238 and Cameor (*P. sativum* subsp. *sativum*) shown to have the slow SCA electromorph (SCA^s^), predominating overwhelmingly in the cultivated pea but occurring in wild peas as well. The derived amino acid sequences had 19 and 18 positively charged amino acid residues in the first and second group, respectively. Full correspondence of the electrophoretic and sequence data, together with the genomic position and the size and content of the protein product, indicate that *Psat0s797g0160* is the *SCA* gene (the latter designation will be used in the text below).

### The related gene *Psat0s797g0240*

The gene *Psat0s797g0240* resides in the pea genome (cultivar Cameor) in about 25 Kb from *Psat0s797g0160*, the two loci have very similar sequence, and the difference in their coding sequences is just 25 nucleotides (8.2 %). Obviously these genes are paralogs originated by tandem duplication of a genome region. The inferred amino acid sequence of the *Psat0s797g0240* polypeptide product has the same length of 101 amino acid sequences and differs (in cv. Cameor) by eight amino acid substitutions from that of *Psat0s797g0160* (their positions are given in parenthesis): ala→ser (14), arg→lys (32), tre→ala (35), arg→his (36), tre→ser (70), val→gln (72), gln→arg (79), asn→his (80). Due to the two latter substitutions this polypeptide has two positively charged residues more than SCA. The mobility of polypeptides of equal length in the involved electrophoretic procedure in acid denaturing conditions is proportional to the number of positively charged residues, so we can expect the Psat0s797g0240 product mobility to be 11% greater than that of SCA^s^. However, the SAA mobility is only 7% greater than that of SCA^s^ (Smirnova et al., 1992). This is close with the mobility of SCA^f^, which is 5% greater than that of SCA^s^ (Smirnova et al., 1992), as expected from their difference for one positively charged residue. Most probably, *Psat0s797g0240* is not the gene coding for SAA but may encode an immunologically related minor protein SA-a2 with electrophoretic mobility 9% greater than in SCA^s^. Electrophoretic monomorphism of both SAA and SA-a2 so far observed (Smirnova et al., 1992) does not allow us to check these options genetically. We attempted resequencing the *Psat0s797g0240* alleles in the same set of accessions as for *SCA* (see above) using primers Ps_SCA2_1F (5’ CACGTGTTCAATAATCTAACGC) and Ps_SCA2_1R (5’ AAGAAAAAGAAACGAGCCATCA) matching the 5’- and 3’ non-coding regions of PUCA011001169.1 (*Pisum sativum* cultivar Gradus No 2 flattened_line_64181, the whole genome shotgun sequence). Possibly, due to the abundance of poly-A and poly-T in the 5’- and 3’non-coding regions, respectively, amplification was not successful for 7 of 20 accessions involved (JI1794, 721, Pe013, P014, P017, VIR5416, WL1238, Cameor), so only 13 accessions were sequenced. The variation of protein products inferred from the obtained sequences was confined to 5 variable amino acid positions and did not affect electrophoretic mobility.

### Protein product of *SCA* and its variation

The SCA protein product is remarkable for its extreme hydrophily and high content of charged (at normal pH) residues (Table 1). Among its 101 amino acid residues in Cameor, 70 are hydrophylic, of which 30 charged, including 18 positively charged (lysine – 10, arginine – 5, histidine – 3), and 12 negatively charged (glutamate – 10, aspartate – 2) residues. (the numbers of residues almost coinciding with their percentages) (Table 1). Interestingly, 11 of 12 negatively charged residues have positively charged nearest neighbour(s) and 4 of 18 positively charged residues have negatively charged neighbour(s). There are tracts of three (glu-lys-glu) and five (lys-lys-glu-glu-arg) charged residues in a row. There are only two proline residues and only one aromatic residue (tyrosine). Such a content and structure, with alternating residues of opposite charge, suggest that in water solution, the SCA molecule has a rigid expanded (linear) structure.

**Table 1.**
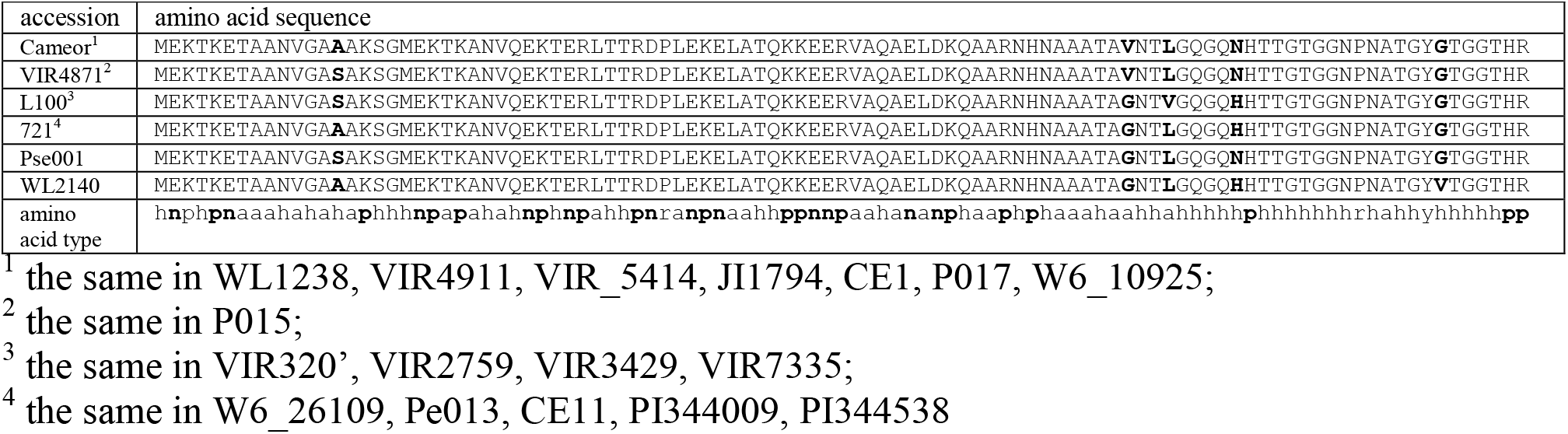
Variants of the deduced amino acid sequences encoded by *SCA* alleles sequenced from pea accessions (polymorphic positions boldfaced). The lowest line shows amino acid types encoded as follows: a – alyphatic, h – hydrophylic, n – negatively charged in neutral conditions, p – positively charged in neutal conditions, r – proline, y – aromatic (tyrosine) (charged types boldfaced)

Five variable amino acid positions were revealed in the SCA product: 14 (ala/ser), 72 (gly/val), 75 (leu/val), 80 (his/asn) and 95 (gly/val) (Fig. 1), the fourth changing the molecule positive charge as discussed above. The *SCA* gene sequences obtained had 11 (3.6 %) variable nucleotide positions.

**Figure 1.**
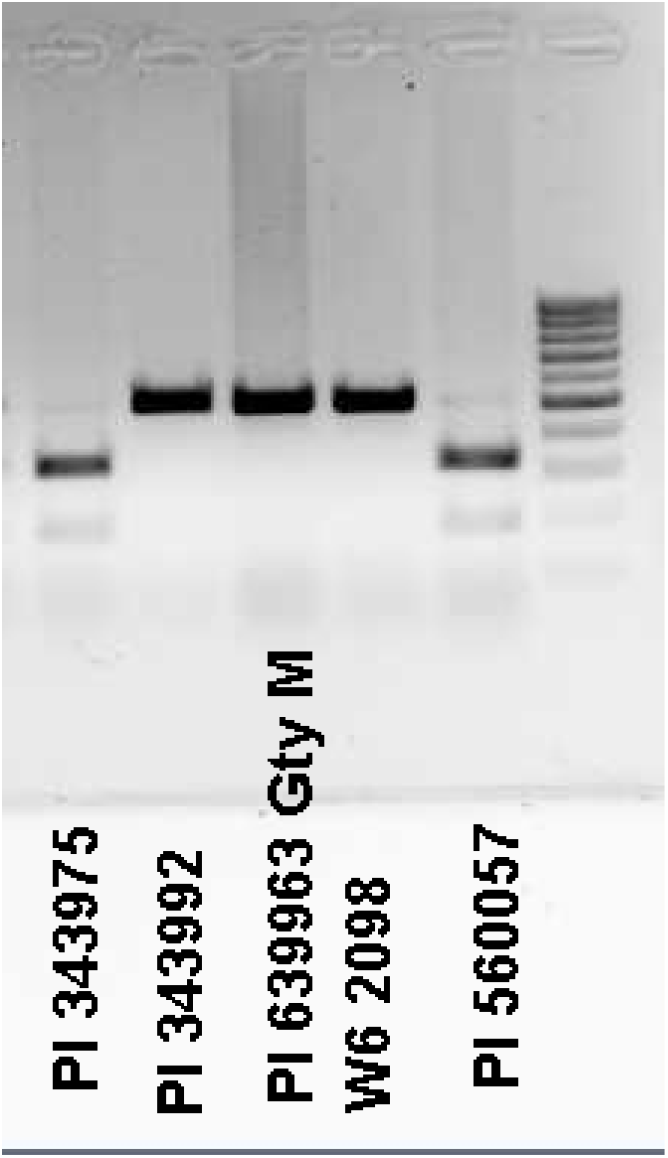
An example of agarose gel electrophoresis of the PCR products obtained from genomic DNA of the indicated pea accessions with primers Ps_SCA1_3F and Ps_SCA1_3R, matching the *SCA* gene adjacent non-coding regions, digested with by *Hind*II restriction endonuclease.

### *SCA* gene as a CAPS marker

The A→C substitution in position 238 of the *SCA* gene creates the recognition site GTCAAC for *Hind*II restriction endonuclease, missing in the rest of the gene. PCR amplification of genomic DNA with Ps_SCA1_3F and Ps_SCA1_3R primers resulted in a product of 512 bp, which was subsequently digested with *Hind*II endonuclease. The allele coding for SCA^f^ was cleaved into two fragments, 326 and 186 bp in size, while amplicon from the SCA^s^ allele remained 512 bp, the difference clearly seen on agarose gel (Fig. 1).

This makes SCA a convenient CAPS (cleaved amplified polymorphic sequence) marker which permits scoring *SCA* allelic state without invoking protein electrophoresis. It should be noted that in spite of great similarity of the coding sequences of SCA and Psat0s797g0240, their adjacent non-coding regions appeared diverged enough to avoid cross-amplification.

## Acknowledmenents

The work was supported by the Russian State Scientific Program № FWNR-2022-0017. Sequencing was carried out in Genomic Core Facility SB RAS, Novosibirsk. Assembly of coding sequences from SRA archives was performed at Siberian Supercomputer Center Core Facility and Computational Facility of Novosibirsk State University. Plants were grown in the greenhouse at the SB RAS Artificial Plant Growing Facility.

## References

Ali S.M., Sharma B., Ambrose M.J. Current status and future strategy in breeding pea to improve resistance to biotic and abiotic stresses. Euphytica.1994;73:115–126. DOI 10.1007/BF00027188

Bogdanova V.S., Shatskaya N.V., Mglinets A.V., Kosterin O.E., Vasiliev G.V. Discordant evolution of organellar genomes in peas (*Pisum* L.). Mol.Phyl.Evol.2021;160:107136. DOI 10.1016/j.ympev.2021.107136

Coyne C.J., McGee R.J., Redden R.J., Ambrose M.J., Furman B.J., Mi-les C.A. Genetic ajustment to changing climates: pea. (Eds S.S. Yadav, R.J. Redden, J.L. Hatfield, H. Lotze-Campen, A.E. Hall). Crop Adaptation to Climate Change Wiley-Blackwell, Oxford, UK, 2011;238–250.

Ford-Lloyd B.V., Schmidt M., Armstrong S.J., Barazani O., Engels J., Hadas R., Hammer K., Kell S.P., Kang D., Khoshbakht K., Li Y., Long C., Lu B.-R., Ma K., Nguyen V.T., Qiu L., Ge S., Wei W., Zhang Z., Maxted N. Crop wild relatives – undervalued, underutilized and under threat? BioScience. 2011;61:559–565. DOI 10.1525/bio.2011.61.7.10

Gorel, F.L., Rozov, S.M., Berdnikov, V.A. Mapping the locus *coch*. Pisum Genetics.1998;30:9–11.

Kalo P., Seres A., Taylor S.A., Jakab J., Kevei Z., Kereszt A., Endre G., Ellis T.H.N., Kiss G.B. Comparative mapping between *Medicago sativa* and *Pisum sativum*. Mol.Genet.Genomics.2004;272(3):235–246. DOI 10.1007/s00438-004-1055-z

Kosterin O.E., Bogdanova V.S. Relationship of wild and cultivated forms of *Pisum* L. as inferred from an analysis of three markers, of the plastid, mitochondrial and nuclear genomes. Genet.Resour.Crop.Evol.2008;55:735–755. DOI 10.1007/s10722-007-9281-y

Kosterin O.E., Bogdanova V.S. Reciprocal compatibility within the genus *Pisum* L. as studied in F_1_ hybrids: 1. Crosses involving *P. sativum* L. subsp. sativum. Genet.Resour.Crop.Evol.2015;62(5):691–709: DOI 10.1007/s10722-014-0189-z

Kosterin O.E., Bogdanova V.S. Reciprocal compatibility within the genus *Pisum* L. as studied in F_1_ hybrids: 3. Crosses involving *P. sativum* L. subs. *elatius* (Bieb.) Aschers. et Graebn. Br. Genet.Res.Crop.Evol.2021;68(6):2565–2590: DOI 10.1007/s10722-021-01151-2

Kosterin O.E., Zaytseva O.O., Bogdanova V.S., Ambrose M. New data on three molecular markers from different cellular genomes in Mediterranean accessions reveal new insights into phylogeography of *Pisum sativum* L. subsp. *elatuis* (Beib.) Schmalh. Genet.Resour.Crop.Evol.2010;57:733–739. DOI 10.1007/s10722-009-9511-6

Kreplak K., Madoui M.A., Cápal P., Novák P., Labadie K., Aubert G., Bayer P.E., Gali K.K., Syme R.A., Main D., Klein A., Bérard A., Vrbová I., Fournier C., d’Agata L., Belser C., Berrabah W., Toegelová H., Milec Z., Vrána J., Lee H., Kougbeadjo A., Térézol M., Huneau C., Turo C.J., Mohellibi N., Neumann P., Falque M., Gallardo K., McGee R., Tar’an B., Bendahmane A., Aury J.M., Batley J., Le Paslier M.C., Ellis N., Warkentin T.D., Coyne C.J., Salse J., Edwards D., Lichtenzveig J., Macas J., Doležel J., Wincker P., Burstin J. A reference genome for pea provides insight into legume genome evolution. Nature Genet.2019;51:1411–1422. DOI 10.1038/s41588-019-0480-1

Maxted N. Ambrose M. 2001. Peas (*Pisum* L.). In: Plant genetic resources of legumes in the Mediterranean (Eds. Maxted N. and S.J. Bennett). Kluwer Academic Publishers, The Netherlands, 2001;181–190.

Maxted N., Kell S.P. Establishment of a global network for the in situ conservation of crop wild relatives: status and needs. FAO Commission on Genetic Resources for Food and Agriculture. Rome, 2009; 266 pp.

Maxted N., Kell S., Ford-Lloyd B., Dulloo E., Toledo Á. Toward the systematic conservation of global crop wild relative diversity. Crop Sci.2012;52:774–785. DOI 10.2135/cropsci2011.08.0415

Panyim, S., Chalkley, R. High resolution in acrylamide gel electrophoresis of histones. Arch.Biochem.Biophys.1969;130:336–346. DOI 10.1016/0003-9861(69)90042-3

Rozov, S.M., Temnykh, S.V., Gorel’, F.L., Berdnikov, V.A. A new version of pea linkage group 5. Pisum Genetics,1993;25:46–51.

Smirnova, O.G., Rozov S.M., Kosterin O.E., Berdnikov V.A. Perchloric acid extractable low-M_r_ albumins SCA and SAA from cotyledons and seed axes of pea (*Pisum sativum* L.). Plant Sci.1992;82:1–13. DOI 10.1016/0168-9452(92)90002-4

Smirnova, O.G., Rozov S.M., Kosterin O.E. Perchloric acid extraction of pea seed proteins: characterisation and inheritance of electrophoretic variants. In: Genetics of Economically Valuable Characteristics of Higher Plants. Institute of Cytology and Genetics os Siberian Branch od the Academy of Sciences of the USSR, Novosibirsk, 1990;158–179 (in Russian).

Smýkal P., Aubert G., Burstin J., Coyne C.J., Ellis N.T., Flavell A.J., Ford R., Hýbl M., Macas I., Neumann P., McPhee K.E., Redden R.J., Rubiales D., Weller J.L., Warkentin T.D. Pea (*Pisum sativum* L.) in the genomic era. Agronomy.2012;274–115. DOI 10.3390/agronomy2020074

Zaytseva O.O., Bogdanova V.S., Kosterin O.E. Phylogenetic reconstruction at the species and intraspecies levels in the genus *Pisum* L. (peas) using a histone H1 gene. Gene.2012;504:192–202. DOI 10.1016/j.gene.2012.05.026

Zaytseva O.O., Gunbin K.V., Mglinets A.V., Kosterin O.E. Divergence and population traits in evolution of the genus *Pisum* L. as reconstructed using genes of two histone H1 subtypes showing different phylogenetic resolution. Gene.2015;556:235–244. DOI 10.1016/j.gene.2014

Zaytseva O.O., Bogdanova V.S., Mglinets A.V., Kosterin O.E. (2017) Refinement of the collection of wild peas (*Pisum* L.) and search for the area of pea domestication with a deletion in the plastidic *psbA-trnH* spacer. Genet.Res.Crop.Evol.2017;64:1417–1430. DOI 10.1007/s10722-016-0446-4

